# Integrative metagenomics and structural bioinformatics identify explainable gut microbial variants associated with Crohn’s disease

**DOI:** 10.64898/2025.12.29.696894

**Authors:** Nadeem Khan, Muhammad Muneeb Nasir, Ubair Aziz, Haseeb Manzoor, Muhammad Faheem Raziq, Zamir Hussain, Ishrat Jabeen, Masood Ur Rehman Kayani

## Abstract

Metagenomics has revealed disease-associated shifts in microbial taxa and functions in inflammatory bowel disease (IBD) patients. However, the role of genomic variation in gut commensals remains poorly understood. Here, we integrated metagenomic profiling, variant calling, and structural bioinformatics to identify disease-associated variants in the gut microbes. Crohn’s disease (CD) and ulcerative colitis (UC) showed significant negative associations with *Bacteroides uniformis*, *Bacteroides vulgatus*, and *Eubacterium rectale*. These bacteria exhibited 190,712 single-nucleotide polymorphisms, including 479 CD-specific and 235 UC-specific variants. Variant prioritization identified a CD-specific Val170Leu substitution in the conserved starch-binding domain of the Starch Utilization System D (SusD) protein in *B. uniformis*. Structural modeling and cyclodextrin docking indicated reduced binding affinity in the mutant, while 200-ns molecular dynamics simulations showed stable ligand retention only in the wild type. These findings suggest that impaired starch metabolism driven by SusD variation may contribute to *B. uniformis* depletion in CD and demonstrate the value of integrating metagenomics with structural analyses to identify functionally relevant microbial variants.

## Background

Inflammatory bowel diseases (IBD), including Crohn’s disease (CD) and ulcerative colitis (UC), represent a growing global public health challenge. In 2017, the worldwide burden of IBD reached approximately 6.8 million cases, with prevalence rising from 79.5 to 84.3 per 100,000 population between 1990 and 2017 [1]. IBD is characterized by chronic, uncontrolled inflammation of the gastrointestinal tract, leading to debilitating symptoms such as abdominal pain, severe diarrhea, rectal bleeding, and intestinal obstruction. While the exact etiology remains incompletely understood, both CD and UC arise from a complex interplay of genetic susceptibility and environmental triggers [2–6].

A critical concern in IBD management is the heightened risk of intestinal malignancies. Due to overlapping histopathological features, misdiagnosis or delayed diagnosis can accelerate progression to colorectal cancer (CRC), anal cancer, intestinal lymphoma, and small bowel adenocarcinoma [7–9]. A global meta-analysis of 116 studies revealed that UC patients exhibit a 3.7% higher prevalence of CRC compared to the general population [10]. Similarly, a population-based study in Manitoba, Canada, reported that individuals with UC and CD face a 2 to 3-fold increased risk of colon cancer [11]. These findings underscore the urgent need for novel molecular insights to improve early detection and precision management of IBD.

Emerging evidence highlights the gut microbiome as a key player in IBD pathogenesis. Dysbiosis marked by shifts in microbial composition and function has been implicated in the distinct pathophysiology of CD and UC [12, 13]. For instance, Raed M. Alsulaiman et al. (2023) identified phenotype-specific differences in genera such as *Phascolarctobacterium*, *Maegamonas*, *Harryflintia*, and *Akkermansia* between UC and CD patients [12]. CD patients also exhibit enriched *Escherichia coli* compared to healthy controls, a trend not observed in UC [14]. Functional studies further reveal disease-associated microbial pathways, including amino acid metabolism, nucleotide biosynthesis, and carbohydrate degradation [15]. Notably, *Bacteroides vulgatus* and its protease proteins were overabundant in UC, suggesting a potential role in mucosal inflammation [16].

Despite these advances, the contribution of microbial genomic variation to IBD pathogenesis remains poorly characterized. Single-nucleotide polymorphisms (SNPs) in microbial genomes may significantly alter gene function and host-microbe interactions, yet comprehensive analyses of these variations in IBD are lacking [17, 18]. We hypothesized that CD and UC are associated with distinct profiles of microbial genomic variants that functionally contribute to disease-specific pathophysiology. To test this hypothesis, we implemented an analytical pipeline **(Figure 1)** to systematically identify and characterize gut microbial genomic variants in CD and UC patients. By combining high-resolution metagenomic analysis with microbial variant calling, we identified phenotype-specific variations and employed structural bioinformatics approaches, including molecular docking and molecular dynamics (MD) simulations, to investigate their potential functional implications. We believe our findings provide new insights into the distinct molecular features of CD and UC, which could inform the development of more precise diagnostic and therapeutic strategies in the future.

**Figure 1:**
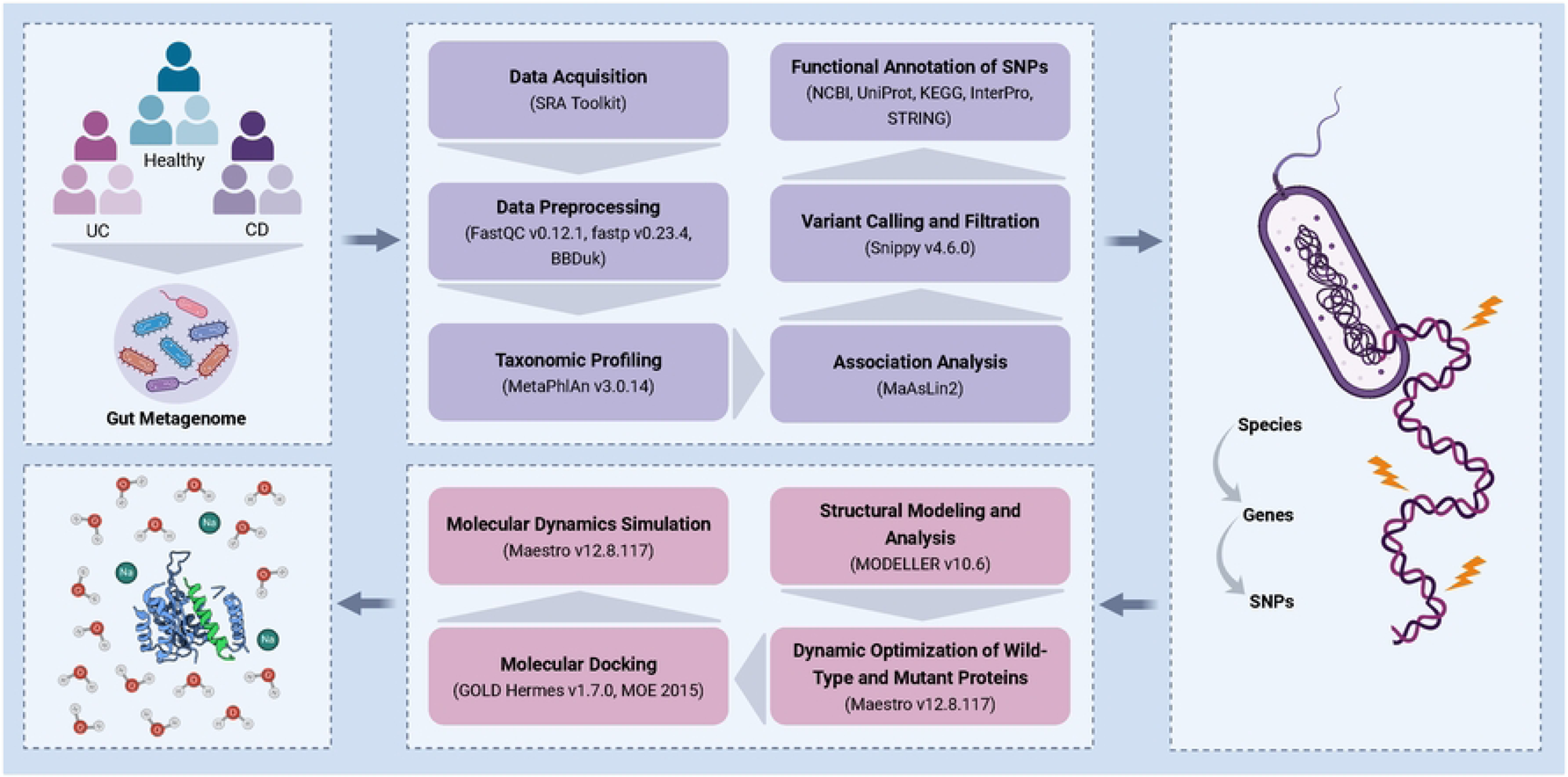
Overview of the Research workflow. The figure shows the complete workflow for identifying gut microbial SNPs specific to CD and UC and associating their effect using structural bioinformatics approaches.

## Materials and methods

### Data collection and preprocessing

We analyzed a total of 150 gut metagenomic samples, comprising CD and UC patients, and healthy controls, obtained from the NCBI SRA database using accession number PRJNA945504 (see **Supplementary Table 1** for detailed sample characteristics) [19]. Read quality was assessed through FastQC (v0.12.1) [20], whereas preprocessing was performed using fastp (v0.23.4) [21]. Preprocessing parameters including: -f 15 (for trimming the first 15 bases of each read), -t 20 (for trimming the last 20 bases of each read), -D (for removing duplicated reads) and -l 70 (for setting the minimum allowed read length to 70) were used. Furthermore, the removal of host reads was performed using BBduk by aligning the reads to the human reference genome (GRCh38) [22] to obtain high-quality reads. The average raw read count was ∼21.24 ± 4.08 Mbps per sample, whereas the post-processed read count was ∼20.46 ± 4.40 Mbps per sample.

### Taxonomic profiling and association analysis

The post-processed reads were taxonomically classified using MetaPhlAn3 (v3.0.14) [23] with the default parameters, except “--ignore_eukaryotes” and “--tax_lev s”. The taxonomic profiling identified 557 bacterial species from ∼30 genera from all the samples **(Supplementary Table 2)**. To identify taxa associated with the three phenotypes, we employed Microbiome Multivariable Associations with Linear Models (MaAsLin2) [24] using parameters; --reference Healthy (setting the healthy group as the baseline for comparisons), --analysis_method LM (linear model framework), --min_prevalence 0.1 (including taxa present in at least 10% of samples), --min_abundance 0.0 and --min_variance 0.0 (retaining all features without abundance or variance filtering), --correction BH (Benjamini–Hochberg procedure for multiple testing correction), and --max_significance 0.05 (FDR threshold of ≤0.05 for statistical significance). Furthermore, significant associations common to CD and UC with prevalence >80% across samples were selected for downstream variant calling.

### Variant calling, filtration, and annotation

To characterize genomic variations in the significantly associated microbial species (**Supplementary Table 3**), we performed variant calling using Snippy (v4.6.0) [25]. This analysis identified multiple variant classes, including SNPs, multiple nucleotide polymorphisms (MNPs), insertions, deletions, and indels, which were further classified as synonymous and nonsynonymous variants based on their predicted functional effects. For downstream analysis, we focused exclusively on nonsynonymous SNPs, as these variants alter amino acid sequences and are more likely to impact protein function.

To identify disease-specific microbial signatures, we implemented a multi-step filtering approach. First, we excluded SNPs commonly present in both healthy controls and diseased (CD/UC) samples, as these likely represent benign polymorphisms of the core microbial genome. This filtering step was crucial to eliminate variants unrelated to disease pathophysiology. Next, we treated SNPs shared between CD and UC cases as baseline microbial genomic variation (background SNPs), as these might reflect general IBD-associated dysbiosis rather than phenotype-specific characteristics. By focusing exclusively on variants unique to either CD or UC, we could identify strain-level microbial differences potentially contributing to each disease’s distinct pathogenesis. Finally, we applied stringent quality thresholds, prioritizing nonsynonymous SNPs with ≥100× sequencing coverage. This depth filter served two critical purposes: (1) ensuring high-confidence variant calls by exceeding typical requirements for microbial variant detection (typically 30-50×), and (2) minimizing false positives that could arise from low-abundance strains or sequencing errors. The resulting high-quality, phenotype-specific SNPs set formed the basis for our downstream functional analyses.

### Functional annotation of SNPs

To functionally characterize the prioritized microbial SNPs in CD and UC, we performed comprehensive annotation of their corresponding proteins using multiple bioinformatics resources. Protein domains and families were analyzed through InterPro, while KEGG facilitated pathway mapping and functional classification. We examined protein-protein interaction networks using STRING and gathered standardized protein annotations from UniProtKB [27–30]. Additionally, we conducted systematic literature searches in PubMed to identify established associations between these proteins and IBD pathogenesis, as well as to evaluate potential functional impacts of the observed mutations.

### Structure modeling and analysis

Through functional annotation and literature curation, we identified a clinically relevant SNP in the *Bacteroides uniformis SusD* gene for detailed analysis, given its established role in polysaccharide utilization and potential to modulate host-microbe interactions. The selected variant encodes a valine-to-leucine substitution at position 170 (V170L) in the SusD protein. We designated the Val170 form as wild-type SusD and the Leu170 variant as mutant SusD. Three-dimensional structures for both protein versions were generated using MODELLER v10.6 through homology-based modeling [30]. The optimal structural template was identified by BLASTp searches against the Protein Data Bank (PDB), which revealed *Bacteroides thetaiotaomicron* SusD (PDB ID: 3CK7; 2.10 Å resolution) as the closest homolog, with ∼25% sequence identity and ∼90% query coverage. This template was subsequently employed to generate high-confidence structural models for downstream comparative analyses. [31, 32]. Ten models each for the wild-type and mutant SusD proteins were generated to evaluate SNP-induced conformational changes. The best models were further refined through energy minimization and MD simulations using Maestro v12.8.117 (Schrödinger Suite), employing the OPLS-2005 force field [33].

### Molecular docking

Following protein modeling and structural refinement, we performed molecular docking to investigate the biological impact of SNP-induced variations on substrate binding. The binding site of the query proteins was selected that includes 10 Å of area surrounding the SNP residue, to precisely evaluate the SNP-induced structural alterations upon ligand interactions. Subsequently, docking simulations were performed using MOE 2015 with default parameters for placement and refinement. The GBVI/WSA dG (Generalized Born Volume Integral / Weighted Surface Area) scoring function was used for both pose generation and refinement, generating 100 ligand conformations for each wild and mutant protein [35]. The mentioned scoring function estimates the free energy of binding (ΔG*_bind_*) through the following empirical equation (1):

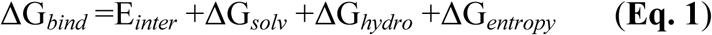

where E₍*_inter_*₎ is the interaction energy (electrostatic and van der Waals), ΔG₍*_solv_*₎ is the desolvation energy from the GBVI model, ΔG₍*_hydro_*₎ represents the hydrophobic contribution estimated via weighted surface area (WSA), and ΔG₍*_entropy_*₎ is the entropy penalty upon ligand binding. This function integrates interaction, solvation, and entropic effects to provide a refined estimate of binding affinity.

### MD simulations

MD simulations were conducted for 200 ns using Maestro v12.8.117 (Schrödinger suite) to assess the stability and conformational changes of the docked complexes over time [34]. The OPLS-2005 force field was employed to model molecular interactions, as it provides a well-validated balance between accuracy and computational efficiency in simulating biological systems [33]. The total potential energy computed for the complexes followed equation (2):

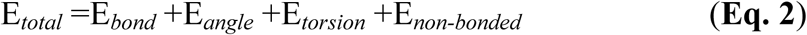

Where E*_bond,_* E*_angle_* and E*_torsion_* represent bonded interactions including bond stretching, angle bending, and torsional rotations, respectively. E*_non-bonded_* accounts for van der Waals and electrostatic interactions between atoms not directly bonded. This comprehensive formulation enables realistic modeling of molecular geometry and energetics in biological systems.

Furthermore, the simulation system was solvated using a TIP3P water model within an orthorhombic box, maintaining a 10 Å buffer around the complex [36]. The system was further neutralized with 0.15 M NaCl, and standard simulation conditions were applied, including a temperature of 300K and pressure of 1 atm. Equilibration was performed using the NVT and NPT ensembles for 100 ps before running the production phase of 200 ns [37]. Parameters such as Root Mean Square Deviation (RMSD), Root Mean Square Fluctuation (RMSF), and hydrogen bonding were calculated to assess the structural stability, flexibility, and interaction patterns of wild type and mutant complexes.

## Results

### Identification of Disease-Associated Microbial Signatures and Functional Variants

To identify microbial taxa, whose abundance patterns distinguish IBD phenotypes from healthy states, we applied MaAsLin2 using linear model (ML) framework. This approach was specifically chosen to quantify the strength of phenotype-specific associations and identify potential protective species showing consistent depletion across both CD and UC. This revealed 190 significantly associated species, with 68 showing CD-specific and 122 showing UC-specific associations (**Supplementary Figure 1**). Among these, three commensal species, *B. uniformis* (CD FDR=3.01×10⁻⁶, coefficient=6.18; UC FDR=2.02×10⁻⁵, coefficient=5.1), *E. rectale* (CD FDR=1.99×10⁻⁷, coefficient=6.14; UC FDR=2.32×10⁻⁵, coefficient=4.37), and *B. vulgatus* (CD FDR=0.00163, coefficient=3.12; UC FDR=0.0421, coefficient=1.97) demonstrated strong positive associations with health (**Figure 2**). Since healthy status was used as the reference, these results indicate significant depletion of these species in both CD and UC patients, suggesting their protective roles in gut homeostasis.

**Figure 2:**
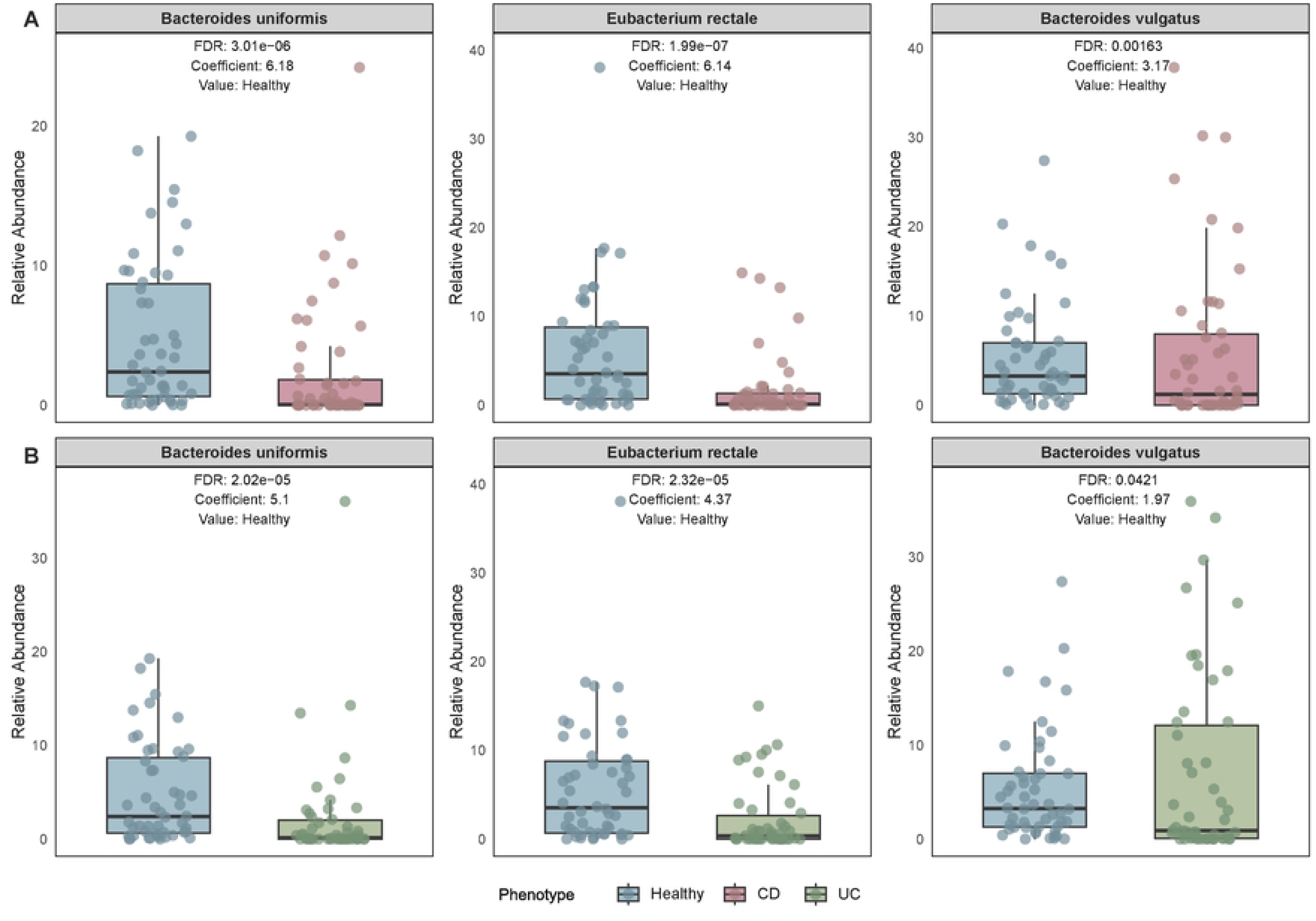
Association analysis of bacterial species with CD and UC. The box plots represent the abundance of bacterial species across the phenotypes, along with significant association coefficients (FDR ≤ 0.05). Positive coefficient values, with healthy as the reference, indicate that the abundance of the mentioned bacterial species is inversely proportional to CD (A) and UC (B), and vice versa.

We hypothesized that strain-level genomic variations in these negatively associated commensals might contribute to their depletion patterns in CD and UC. To investigate this possibility, we performed comprehensive variant analysis of these three species, identifying >1.4M total genomic variations across all samples, including 269,844 non-synonymous mutations (**Supplementary Table 4, Supplementary Figure 2**). The SNP distribution patterns revealed intriguing phenotype-specific differences: while *B. uniformis* showed progressive SNP reduction from healthy (22,290) to CD (18,009) to UC (11,837), *B. vulgatus* exhibited an opposite trend with CD-associated increase (healthy: 34,912; CD: 40,361; UC: 22,718). From these, we focused on high-confidence variants unique to each phenotype (**Table 1**), ultimately prioritizing 714 SNPs with sequencing depth >300× (CD:479; UC:235) (**Supplementary Figure 3**, **Supplementary Table 5**) for downstream functional analysis. This stringent selection was based on the premise that these strain-specific variations might mechanistically explain the observed depletion patterns of beneficial microbes in IBD pathogenesis.

**Table 1.** Comparative summary of synonymous and nonsynonymous variants across bacterial species associated with CD, UC, and healthy phenotypes.

| **Variants** | ***Bacteroides***<br>***uniformis*** | ***Bacteroides***<br>***vulgatus*** | ***Eubacterium***<br>***rectale*** |
| --- | --- | --- | --- |
| Synonymous variants | 303,996 | 484,716 | 365,780 |
| Non-synonymous variants | 80,184 | 131,324 | 58,335 |
| SNPs with non-synonymous variants | 52,136 | 97,991 | 40,585 |
| Unique SNPs in CD | 2455 | 2030 | 1069 |
| Unique SNPs in UC | 1079 | 1754 | 737 |
| Unique SNPs in healthy | 1463 | 1300 | 2721 |

### SNP in SusD Implicates Disruption of the Starch Utilization pathway of *B. uniformis* in CD patients

To identify functionally relevant variants, we implemented a stringent filtering approach that selected for nonsynonymous SNPs within evolutionarily conserved protein domains. This analysis yielded 244 high-priority SNPs (164 CD-associated and 80 UC-associated variants). The biological relevance of these variants was further supported by existing literature demonstrating that impairment of bacterial polysaccharide utilization systems particularly those involved in SCFA production can disrupt mucosal homeostasis and promote inflammatory responses. To functionally characterize the 244 prioritized variants, we performed comprehensive annotation of the affected proteins and filtered for those involved in carbohydrate metabolism using relevant keywords (polysaccharide, glucan, starch, binding, and metabolism). This analysis identified 41 proteins functionally linked to polysaccharide utilization pathways, including 10 in *B. uniformis*, 15 in *B. vulgatus*, and 16 in *E. rectale* (**Supplementary Table 6**).

Notably, several proteins identified in this subset, including phosphopyruvate hydratase (eno), glucose-1-phosphate adenylyltransferase subunit GlgD (glgD), L-arabinose isomerase (araA), glycerol kinase (glpK), and 2,3-bisphosphoglycerate-independent phosphoglycerate mutase (gpmI), play established roles in carbohydrate metabolism. Among these, a nonsynonymous SNP in the SusD protein, a key component of the starch utilization system (Sus) emerged as particularly relevant. This SNP, with a high sequencing depth (562X), involved a valine-to-leucine substitution at the 170th amino acid position (Val170Leu), specifically in *B. uniformis* from CD patients. Furthermore, the SNP was located within the conserved domain of the SusD protein (residues 89–230), which is essential for its starch-binding function. Network analysis further underscored the functional importance of SusD, revealing direct associations with other core proteins of the starch utilization system (SusB, SusC, SusE, and SusF), with positive co-expression scores of 0.067, 0.050, 0.073, and 0.103, respectively, each predicted with a confidence level exceeding 70% **(Supplementary Figure 4)**.

### Molecular Modeling Revealed Local Structural Perturbations in Mutant SusD

We hypothesized that the Val170Leu mutation in *Bacteroides uniformis* SusD would induce local structural alterations affecting its starch-binding capacity, despite maintaining overall protein stability. To test this, we modeled structures of wild-type (Val170) and mutant (Leu170) SusD followed by MD simulations. Comprehensive sequence alignment revealed substantial regions of discontinuity between template and target sequences (**Supplementary Figure 5**). Most critically, the mutation locus and its flanking residues (Ile169, Thr171, Ser172, Tyr173, Glu174, Gln175, Pro176, Gln177, Asp178, Glu179, and Val180) resided within an extensive unstructured loop region completely absent from the template structure. This structural gap encompassed eleven consecutive residues proximal to the polymorphic site, including several residues predicted to participate in substrate coordination based on homologous systems.

Targeted loop refinement was performed using MODELLER v10.6 with the ‘very_slow’ refinement protocol (incorporating slow, iterative optimization steps) to specifically address the unresolved region. This high-stringency approach successfully resolved the backbone conformation for four consecutive residues (Ile169, Val170, Thr171, Ser172) in the wild-type structure and three residues (Ile169, Val170, Thr171) in the mutant variant, thereby enabling comparative structural analysis of this functionally critical region.

Simulation studies revealed critical differences in protein behavior between wild-type and mutant SusD variants. Global stability analysis showed comparable RMSD profiles for both structures, maintaining an average deviation of ∼6 Å after 80 ns with similar fluctuation patterns throughout the 200 ns simulation (**Supplementary Figure 6A**). RMSF analysis demonstrated that while the overall structure remained stable, the mutant SusD exhibited increased flexibility at specific residues, including Ser58, Glu59, Asp87, Asn209, Met211, Leu346, Ala347, and Arg466. These residues showed substantially greater fluctuations in the mutant variant (average ∼5 Å) compared to the wild type (∼2 Å) (**Supplementary Figure 6B**). The mutation-induced flexibility changes were particularly pronounced in regions surrounding the substrate-binding site, suggesting potential functional consequences for starch binding and processing.

We hypothesized that the Val170Leu substitution would induce localized conformational changes in SusD’s starch-binding domain while preserving global protein architecture. Supporting this prediction, comparative analysis of the 200 ns simulation endpoints revealed significant local reorganization around the mutation site (Thr171-Asp178; **Figure 3**), despite maintained global structural integrity. Quantitative superposition with the wild-type structure identified three distinct perturbation clusters: (1) a severely displaced region centered on Glu174 (21.0 Å deviation), (2) intermediate displacements at Pro176 (13.1 Å) and Gln177 (10.3 Å), and (3) moderate but functionally relevant shifts at the mutation site (Leu170: 6.0 Å) and adjacent residues (average regional deviation: 12.0 Å). These findings demonstrate that the Val170Leu mutation creates a cascade of structural perturbations radiating through the binding domain, with magnitude inversely proportional to distance from the mutation site, a pattern consistent with allosteric modulation of substrate recognition geometry.

**Figure 3:**
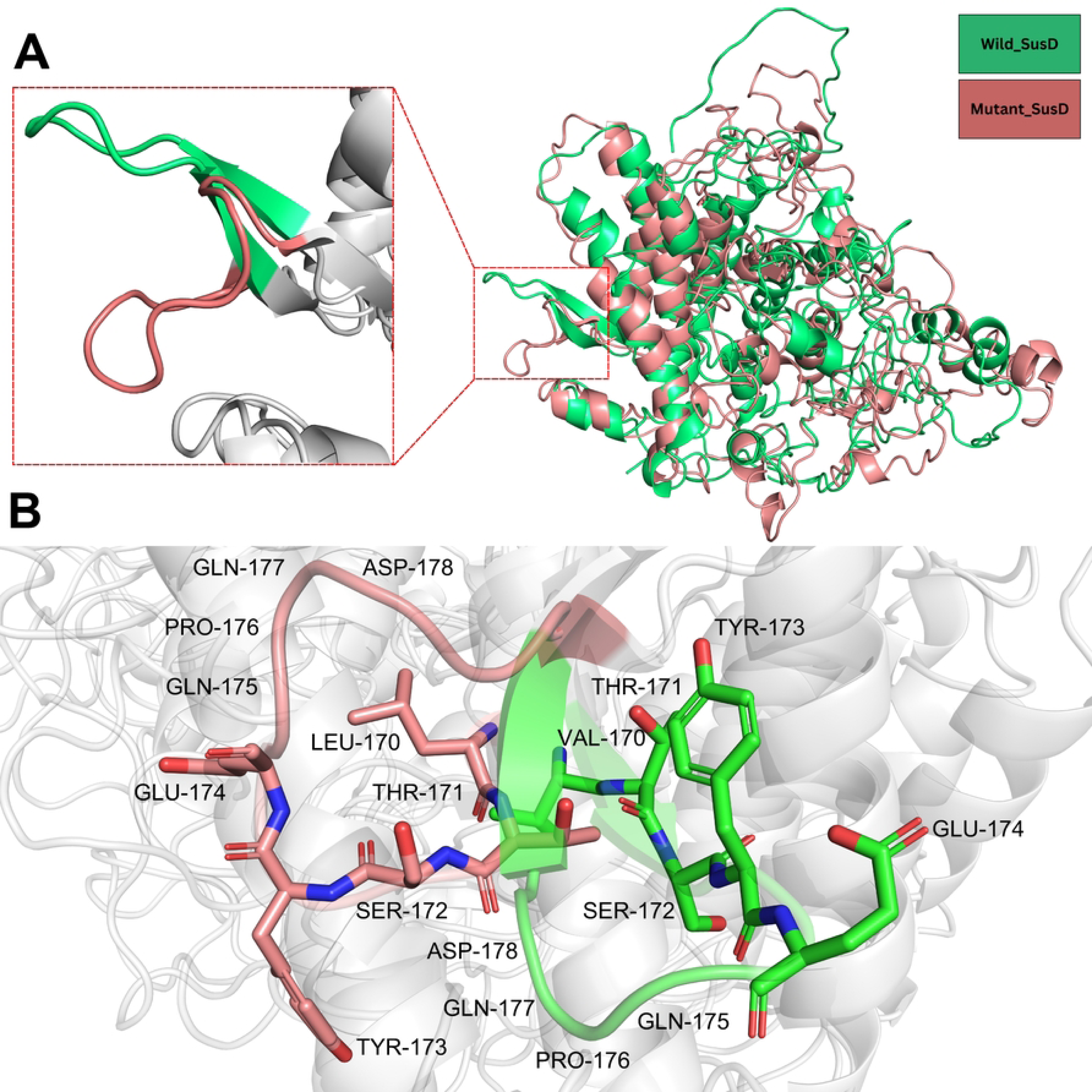
Structural superimposition of wild and mutant SusD. (A) Superimposed structures of wild and mutant SusD, where the region, encompassing the variation (Val170Leu) along with its neighboring residues, showed a major structural deviation. (B) Conformational deviations of each residue present along the variation site, demonstrating the extent of deviation between each residue of wild and mutant SusD.

### Docking Analysis Indicates Weakened Cyclodextrin Binding in Mutant SusD

To evaluate whether the observed structural perturbations affected ligand binding, we performed molecular docking of cyclodextrin, an oligosaccharide mimicking starch into the binding sites of both wild-type and mutant SusD. Cyclodextrin (**Supplementary Figure 7**), originally co-crystallized with the template protein (PDB ID: 3CK7), was docked into the binding pocket of each version of protein, generating 100 conformations per complex. Comparative analysis revealed significant differences in binding energies and interaction networks between the wild-type and mutant complexes. The wild-type SusD exhibited stronger cyclodextrin binding, with an average binding energy of −6.274 kcal/mol across all conformations compared to −5.496 kcal/mol for the mutant. This trend was particularly evident in the top-scoring poses, where the wild-type complex achieved a binding energy of −9.125 kcal/mol versus −6.092 kcal/mol for the mutant (**Supplementary Figure 8**).

Detailed examination of binding interactions showed that the wild-type SusD maintained a more extensive network of stabilizing contacts with cyclodextrin. Specifically, five residues (Ser123, Ser126, Asp131, Ser172, and Asp178) participated in hydrogen bonding with the ligand, while only three residues (Asn124, Glu174, and Gln175) were involved in the mutant complex (**Figure 4A and C**). Similarly, non-bonded interactions were more numerous in the wild-type complex, involving ten residues (Leu122, Leu127, Ile130, Ser134, Arg153, Val170, Thr171, Pro176, Gln177, and Val180) compared to seven residues (Leu116, Val120, Tyr173, Pro176, Lys523, Asp526, and Leu527) in the mutant (**Supplementary Figure 9A and C**).

**Figure 4:**
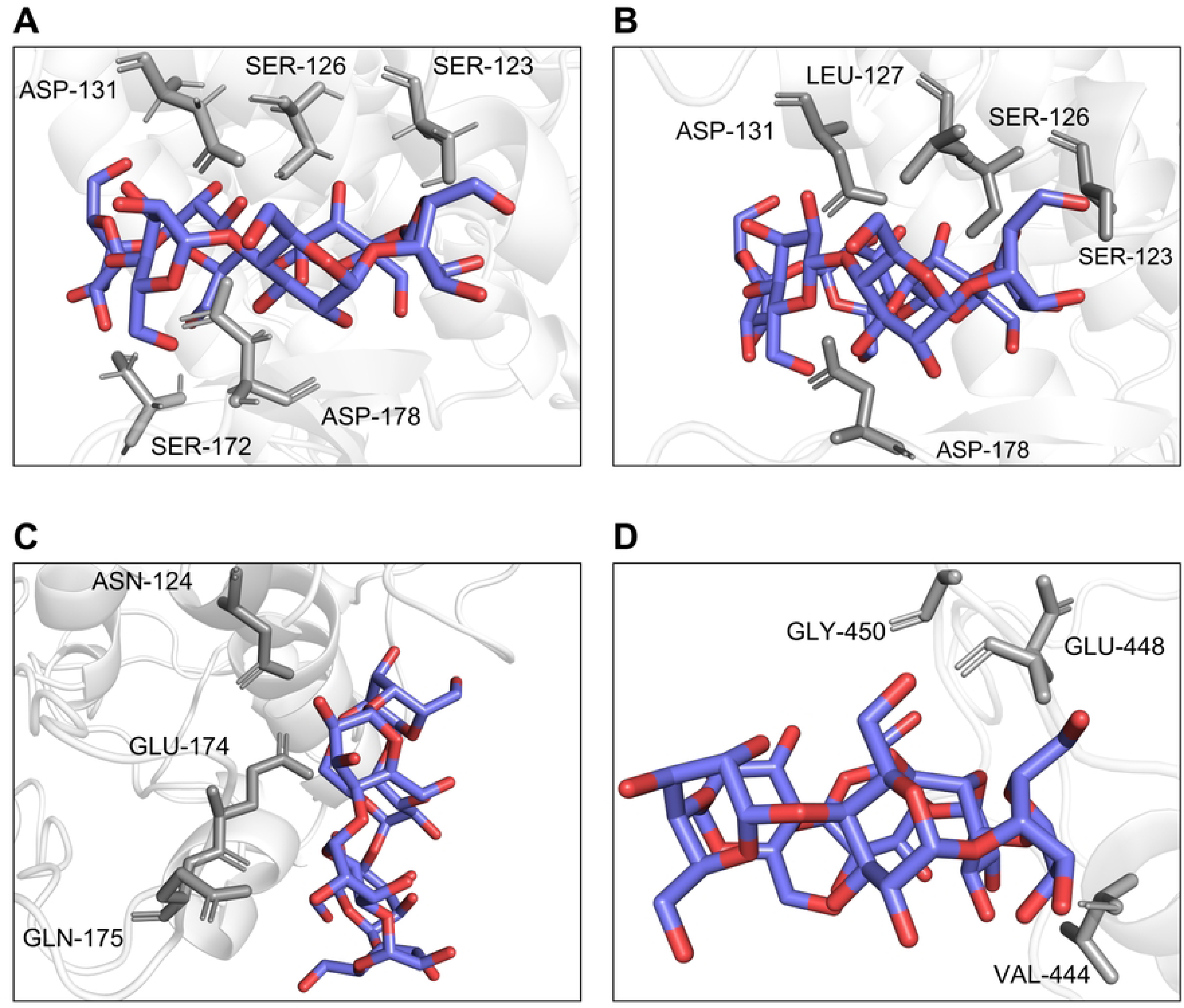
Residues of wild and mutant SusD making hydrogen bonds with cyclodextrin before and after simulation. (A) Residues of wild SusD involved in hydrogen bond formation with cyclodextrin after docking, while (B) Interacting residues following molecular dynamics simulation. (C) Residues of mutant SusD contributing to hydrogen bonding with cyclodextrin after initial docking. (D) Corresponding interactions after simulation.

### Comparative MD Simulations Highlight Impaired Ligand Retention in Mutant SusD

Building upon our previous findings that the Val170Leu mutation induces local structural perturbations in SusD (average deviation of ∼12 Å around residues 170-178), we hypothesized that these conformational changes would specifically disrupt cyclodextrin binding stability.

Molecular dynamics simulations of the docked complexes confirmed this hypothesis, revealing marked differences in ligand interaction patterns between wild-type and mutant forms. The wild-type complex maintained structural integrity throughout the 200 ns simulation, with cyclodextrin preserving both its binding pose and key interactions observed in docking studies. Hydrogen bonds with Ser123, Ser126, Leu127, Asp131, and Asp178 remained stable (**Figure 4B**), while non-bonded contacts with ten binding site residues persisted (**Supplementary Figure 9B**). This stability contrasted sharply with the mutant complex, where cyclodextrin progressively dissociated from its original position - particularly from the mutation-affected region (residues 170-178) that showed the greatest structural deviations in our earlier analysis. The displaced ligand formed alternative interactions with Val444, Glu448, and Gly450 (**Figure 4D**), while losing contact with original binding site residues. Non-bonded interactions similarly shifted to Lys443, Arg445, Asn446, Gly447, Ser451, Leu452, and Glu453 (**Supplementary Figure 9D**).

Comparative RMSD analysis revealed distinct structural stabilization patterns between the wild-type and mutant SusD-cyclodextrin complexes during the 200 ns simulation **(Figure 5A)**. The wild-type complex underwent rapid conformational adjustments during the initial 20 ns, as evidenced by an RMSD increase to ∼5.5-6 Å, after which it maintained stable structural equilibration with minimal fluctuations. In contrast, the mutant complex displayed slower structural reorganization, gradually increasing to ∼5 Å RMSD over 150 ns before achieving relative stability. RMSD analysis of the docked cyclodextrin revealed distinct behaviors: within the mutant complex, it showed initial fluctuations before stabilizing at ∼80 Å displacement, while the wild type reached ∼18 Å RMSD within 45 ns and remained stable (**Supplementary Figure 10**). RMSF analysis (**Figure 5B**) demonstrated greater flexibility (∼5.2 Å) in the mutant’s Thr171-Asp178 region and alternative binding site Val444-Glu453 (∼5.6 Å) compared to the wild-type, consistent with the observed binding site displacement and local structural perturbations.

**Figure 5:**
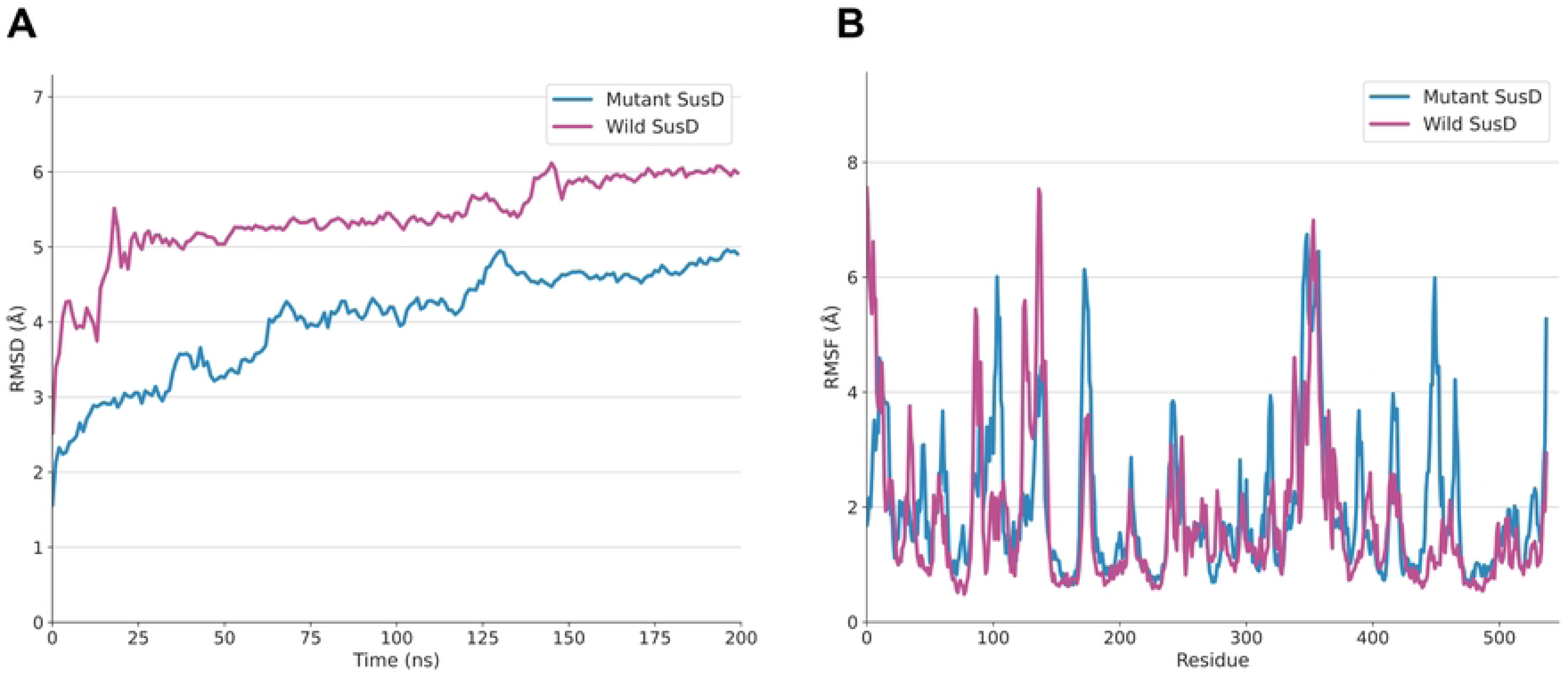
Molecular dynamics simulation of docked complexes. (A) Comparative RMSD of docked wild and mutant SusD along a 200 ns simulation. Both the docked proteins initially deviate from their initial conformation at the beginning of the simulation and stabilize (minimal conformational changes) after the 150th time point of the simulation. (B) Comparative conformational fluctuations of each residue of both docked complexes along the simulation run, where multiple residues of the mutant complex showed notable conformational changes compared to the wild complex, demonstrating the variable interaction of cyclodextrin to multiple residues of the mutant complex at variable locations.

Lastly, we hypothesized that the Val170Leu mutation would disrupt the stable hydrogen bonding network required for productive cyclodextrin binding in SusD. Supporting this prediction, our hydrogen bond analysis revealed markedly different interaction profiles between wild-type and mutant complexes. The wild-type SusD maintained persistent hydrogen bonds with Thr171, Ser172, Tyr173, and Glu174 throughout the simulation (**Supplementary Figure 11A**), preserving the initial binding mode. In striking contrast, the mutant complex failed to sustain these native interactions, instead forming new hydrogen bonds with Val444-Glu453 residues (**Supplementary Figure 11B**), a clear shift from the original binding site. Between 50-150 ns, the mutant complex established consistent but alternative interactions with Glu448, Asp449, Gly450, Ser451, Leu452, and Glu453. Time-resolved conformational analysis (**Figure 6**) visually confirmed these differences, with cyclodextrin maintaining stable positioning in the wild-type complex while progressively dissociating in the mutant. These results demonstrate that the Val170Leu mutation not only alters local protein structure but also critically disrupts the hydrogen bonding network essential for proper substrate recognition and binding.

**Figure 6:**
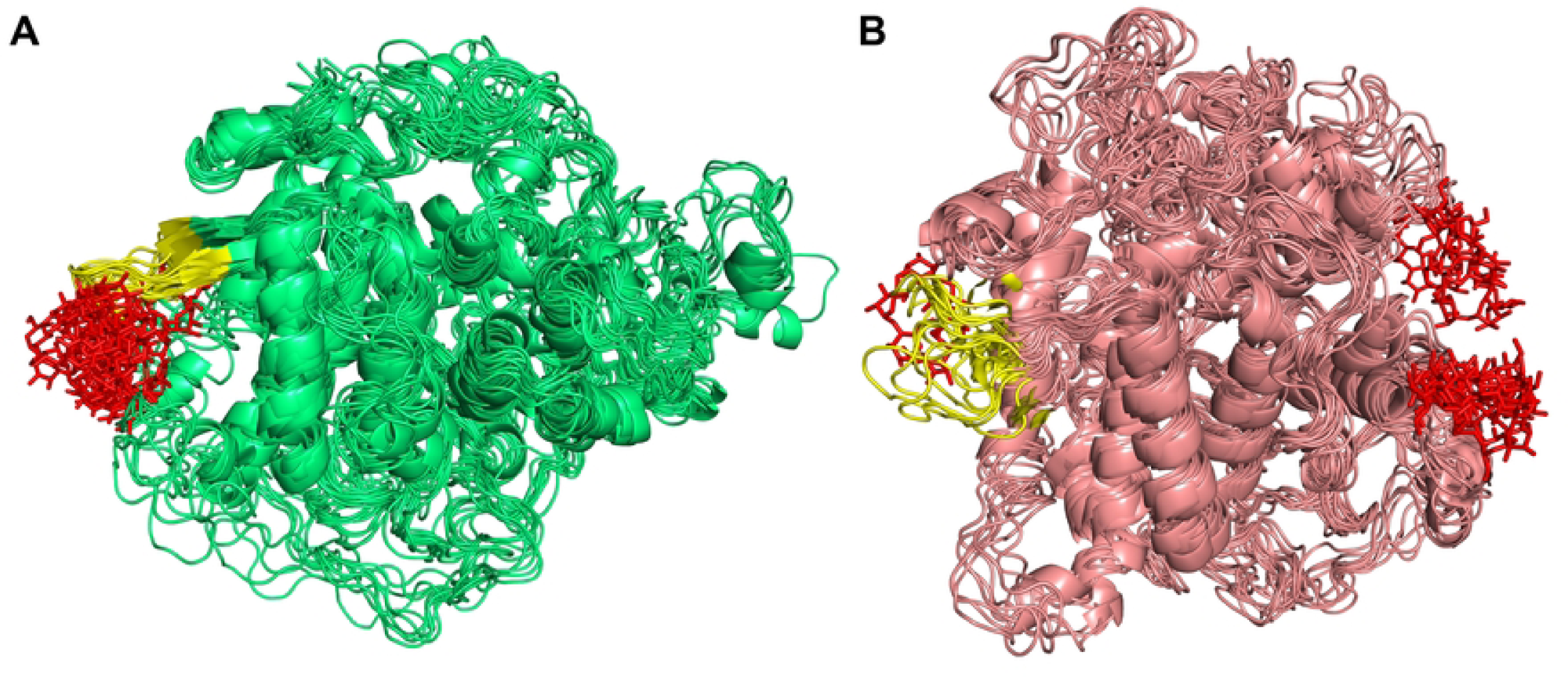
Comparative binding conformations of cyclodextrin at different time points of the MD simulation. Binding conformations of cyclodextrin with wild and mutant SusD at ten evenly spaced time points over the total simulation period (200 ns) were selected. (A) Cyclodextrin exhibited overall stable binding within the chosen docked site (yellow) of wild-type SusD (green). (B) In the mutant SusD (pink), cyclodextrin displayed unstable binding, characterized by notable positional shifts and interactions with other regions (detailed in the main text), highlighting the effect of mutations compared to the wild-type structure.

## Discussion

The chronic inflammation characterizing IBD arises from complex host-microbe interactions that remain incompletely understood. While previous studies have identified shifts in microbial abundance in CD and UC patients, our work provides unprecedented mechanistic insight by characterizing strain-level genomic variations and their structural-functional consequences in key commensal species. Three critical findings emerge from our study: (1) the identification of protective species (*B. uniformis*, *B. vulgatus*, *E. rectale*) showing consistent depletion in both CD and UC; (2) the discovery of phenotype-specific genomic variants in these species; and (3) the structural and functional characterization of a disease-associated SNP in *B. uniformis* SusD protein that impairs starch binding, a potential novel mechanism contributing to pathogenesis in CD.

Our taxonomic profiling and subsequent association analysis are consistent with previous reports of *Bacteroides* depletion in IBD [38, 39]. Furthermore, previous studies have also demonstrated that *B. uniformis, B. vulgatus,* and *E. rectale* exhibit a significant positive association with healthy controls compared to CD and UC phenotypes [39], thus indicating their positive effect on health. The protective role of *B. uniformis* has been experimentally validated in murine models, where administration of this commensal bacterium maintained intestinal homeostasis without inducing adverse histological effects. [45, 46]. Similarly, the abundant nature of *E. rectale* and *B. vulgatus* in the normal human gut compared to CD and UC, and their role in alleviating colitis, has also been reported previously [41, 43].

While the differential abundance patterns of these three bacteria in CD and UC patients provide important ecological insights, these observations alone cannot explain the mechanistic basis of disease pathogenesis. To bridge this knowledge gap, we investigated whether strain-level genomic variations in these commensals might contribute to their depletion and functional impairment in CD and UC. We specifically hypothesized that disease-associated genetic variants in these protective species would be enriched in functional domains of proteins critical for host-microbe interactions. Our analysis revealed substantial genetic divergence, with *E. rectale* exhibiting the highest number of variations, followed by *B. uniformis* and *B. vulgatus.* Among these, a particularly noteworthy finding was a high-confidence, non-synonymous SNP (Val170Leu) located within the functional starch-binding domain of *B. uniformis* SusD - a key component of the *Bacteroides* starch utilization system [38, 50]. This mutation, exclusively identified in CD patients with high sequencing depth (562×), represents a strong candidate for functional impairment, as domain-localized nonsynonymous SNPs are most likely to disrupt protein-substrate interactions critical for commensal fitness and host benefit.

Starch serves as a critical nutritional substrate for *Bacteroides*, functioning as both their primary energy source and a fundamental requirement for maintaining metabolic homeostasis and ecological fitness in the gut environment [45]. Bacteroides-mediated starch metabolism plays a central role in gut microbial cross-feeding, generating metabolic byproducts that serve as key nutrients for other commensal bacteria. This metabolic cooperation underpins critical symbiotic relationships within the gut ecosystem [46]. The starch utilization system (Sus) in *Bacteroides* species comprises a highly conserved seven-protein complex (SusA-G) that orchestrates the efficient breakdown of starch into metabolically useful oligosaccharides and monosaccharides [47–49]. The SusD, SusE, and SusF proteins localized on the outer membrane of *Bacteroides* facilitate starch binding and capture, while SusG mediates its initial hydrolysis into oligosaccharides. These oligosaccharides are then transported across the outer membrane via SusC into the periplasm, where SusA and SusB further degrade them into glucose monomers for cellular uptake and metabolism [32, 50–53]. The complete degradation of starch by gut microbes yields short-chain fatty acids (SCFAs) as primary metabolic end-products, which serve as crucial signaling molecules and energy substrates for both microbial communities and the host. These SCFAs particularly acetate, propionate, and butyrate orchestrate key physiological processes including immune modulation, epithelial barrier maintenance, and energy homeostasis through specific receptor-mediated pathways [54]. SCFAs play a pivotal role in immunomodulation, with *Bacteroides*-derived fermentation products serving dual functions as both the primary energy source for colonocytes and key regulators of inflammatory responses. Through inhibition of histone deacetylases (HDACs) and suppression of NF-κB signaling pathways, these microbial metabolites downregulate proinflammatory cytokine production, thereby maintaining intestinal immune homeostasis and preventing aberrant inflammation [45].

SusD serves as the main starch-binding protein in the *Bacteroides* starch utilization system, playing an essential role in substrate capture within the gut environment. This critical function was experimentally validated by Shipman et al. (2000), who demonstrated that domain-specific mutations in SusD of *B. thetaiotaomicron* completely abolished starch binding capacity, while analogous mutations in other starch-binding proteins maintained partial activity [50]. Our identification of a functional domain mutation (Val170Leu) in the SusD protein of *B. uniformis* from CD patients provides a plausible mechanistic link between microbial genetic variation and intestinal inflammation. We hypothesize this mutation impairs starch binding and disrupts the starch utilization system in *B. uniformis*, reducing glucose and SCFA production. This may explain *B. uniformis* depletion in CD patients, while SCFA deficiency could contribute to disease through: (1) diminished colonocytes energy, (2) dysregulated immune responses, and (3) compromised epithelial integrity [55–58].

The structural perturbations induced by the Val170Leu mutation provide compelling mechanistic support for our hypothesis of impaired starch utilization in CD-associated *B. uniformis*. The persistent conformational changes observed in the binding domain (residues Thr171-Asp178), evidenced by both static structural comparisons and dynamic simulations (∼5.2 Å RMSF increase), suggest a substantial reorganization of the starch-binding interface. While global protein folding remained stable (as indicated by comparable overall RMSD), the localized destabilization of this critical functional region would likely compromise several aspects of substrate interaction: (1) initial starch recognition by SusD, (2) productive positioning for SusG-mediated hydrolysis, and (3) efficient transfer to downstream Sus components. These observations align with established structure-function relationships in starch-binding proteins, where even minor perturbations in binding regions can dramatically reduce substrate affinity [59]. The mutation’s location within SusD’s conserved domain (residues 89-230) further underscores its likely functional significance, as this domain has been evolutionarily optimized for polysaccharide interactions [32]. Notably, the affected residues (Thr171-Asp178) constitute a critical segment that, in homologous systems, directly participates in oligosaccharide coordination through both hydrogen bonding and hydrophobic contacts. This structural evidence, combined with our docking results showing reduced cyclodextrin affinity, strongly suggests that the Val170Leu variant could be a loss-of-function mutation with potential consequences for *B. uniformis* ecology and host-microbe interactions in CD.

Similarly, site-specific docking supports our hypothesis, demonstrating that fewer residues participate in cyclodextrin binding in mutant SusD compared to wild type, resulting in reduced binding affinity. The comparative binding energies, lacking absolute reference values, prompted validation through 200ns simulation studies [60]. MD simulations revealed distinct dynamic behavior of cyclodextrin between the mutant and wild-type SusD complexes. While cyclodextrin maintained stable binding throughout the simulation in the wild-type complex, the mutant complex exhibited early conformational fluctuations followed by stabilization at a significantly displaced position. These observations demonstrate how structural perturbations affect ligand binding and strongly support our hypothesis of mutation-induced disruption in substrate recognition.

The hydrogen bonding patterns provide critical molecular-level insights into the mutation’s functional consequences. The wild-type complex maintained stable interactions with residues 170-174 throughout simulations, explaining its consistent binding pose. In striking contrast, the mutant’s inability to sustain these interactions, instead forming transient bonds before relocating to residues 443-490 - reveals fundamental defects in substrate recognition.

While these alternative interaction sites reside in α-helical regions (Koropatkin et al., 2008), their apparent inability to support productive starch binding suggests the Val170Leu mutation forces compensatory, but functionally inferior, binding modes. This observation aligns with our structural data showing local domain destabilization, collectively demonstrating how single-residue changes can propagate to disrupt entire functional networks in microbial nutrient acquisition systems. [32].

While our findings provide mechanistic insights into how microbial genomic variations may contribute to CD pathogenesis, several limitations warrant consideration. First, although we identified functionally disruptive variants in *B. uniformis* SusD, we did not comprehensively investigate parallel variations in the other two protective taxa (*B. vulgatus* and *E. rectale*), which may harbor equally or more significant disease associations. Second, our computational predictions require validation in independent IBD cohorts to confirm the diagnostic specificity of the Val170Leu mutation. Third, while our structural models strongly suggest impaired starch binding, experimental validation through *in vitro* affinity measurements and *in vivo* colonization assays remains essential. Finally, the limited template similarity (24.9% identity) for homology modeling, particularly in unresolved substrate-binding loops, introduces uncertainty in local conformational predictions. Addressing these limitations through multi-cohort validation, functional assays, and cryo-EM structural studies will strengthen the translational potential of these findings.

## Conclusions

Our integrated analysis identifies the Val170Leu mutation in *B. uniformis* SusD as a potential CD-specific biomarker. By combining high-resolution metagenomic analysis, microbial variant calling, and structural bioinformatics, we demonstrate how this variant disrupts starch-binding functionality a defect that may contribute to inflammation through impaired SCFA production. The consistent computational evidence (reduced binding affinity, ligand displacement, and altered interaction networks) strongly implicates this mutation in CD pathogenesis. While requiring experimental validation, these findings establish a framework for targeting microbial genomic variants in precision IBD management, moving beyond taxonomic profiling to mechanistic, strain-level interventions.

## Declarations

### Ethics Statement and Patient Consent

Not applicable.

### Clinical Trial Number

Not applicable.

### Authors’ contributions

NK, MMN, UA, HM, and MFR performed the analysis. NK prepared the manuscript. MRK conceived the idea. IJ supervised the structural bioinformatics analysis, and ZH guided the statistical analysis. MRK supervised the study and revised the manuscript. All authors reviewed and approved the final manuscript.

## Acknowledgments

We thank the members of the Metagenomics Discovery Lab for their continuous feedback and support in this research. We also thank the SINES Supercomputing Lab for providing the computational resources.

